# Neurodevelopment of Corticostriatal Circuits and Risk for Alcohol Use During the Transition from Adolescence to Adulthood

**DOI:** 10.64898/2026.07.15.738690

**Authors:** Daniel J. Petrie, Ashley C. Parr, Finnegan J. Calabro, Will Foran, Sandra A. Brown, Susan Tapert, Kate Nooner, Douglas Fitzgerald, Duncan Clark, Beatriz Luna

## Abstract

Adolescence and early adulthood are marked by rapid neurobehavioral development in reward and decision-making processes, coinciding with the initiation and escalation of alcohol use. While adolescent and young adulthood alcohol initiation is not atypical, adult trajectories diverge: some individuals reduce or discontinue use, whereas others escalate to more frequent or problematic patterns leading to substance use disorders. Corticostriatal circuits, including the ventral striatum (nucleus accumbens; NAcc) and dorsal striatum (caudate and putamen), support reward processing, goal-directed behavior, and habit formation, and are thought to contribute to distinct stages of alcohol use. Yet, how the normative maturation of these circuits relates to alcohol initiation and the transition to habitual consumption remains unclear. We used data from the National Consortium on Alcohol and NeuroDevelopment in Adolescence and Adulthood (NCANDA-A) cohort (822 participants, baseline ages 12 – 22 years old, 1 – 9 visits per participant, 4,356 total visits), a large multisite longitudinal neuroimaging sample spanning adolescence to young adulthood. We observed that rsfMRI functional connectivity (FC) patterns varied systematically across striatal subdivisions: NAcc FC followed an inverted U-shaped trajectory, peaking during adolescence; while caudate and putamen FC showed monotonic decreases with age. Adolescent peak NAcc connectivity was associated with alcohol use initiation, while a lack of normative decrease in putamen connectivity was linked to more frequent alcohol use in adulthood. Together, results suggest that the maturation of reward processing circuitry may support alcohol initiation, while a lack of habit system specialization may contribute to continued alcohol use, with potential implications for the timing of interventions aimed at limiting at-risk drinking.

## Introduction

Adolescence and early adulthood are characterized by heightened neuroplasticity of cognitive and reward systems in parallel with dynamic behavioral change, during which neural circuits underlying motivation, learning, and decision-making mature rapidly (1). This developmental window coincides with the initiation of alcohol use, which may be driven by a normative increase in sensation seeking (2). However, alcohol use developmental trajectories diverge: many individuals reduce or discontinue use as they mature, whereas others escalate to more frequent or problematic patterns (3). This transition from experimentation to sustained use represents a critical but understudied developmental shift. Early initiation of alcohol use is a well-established risk factor for later substance-related problems (4), yet the neurodevelopmental mechanisms linking normative brain maturation to the onset, escalation, or remission of use remain poorly understood. Identifying neural mechanisms that predict both initiation and maintenance of alcohol use is critical to informing prevention and early intervention strategies during this sensitive developmental period.

Corticostriatal circuits play a central role in motivational and behavioral control, supporting reward-driven, goal-directed, and habitual behaviors (5). Converging evidence suggests that these circuits contribute to different stages of alcohol and substance use. The ventral striatum (nucleus accumbens; NAcc), has been linked to reward sensitivity, risk-taking, and early substance use, and heightened NAcc activation predicts subsequent initiation during adolescence (6,7). In contrast, dorsal striatal regions (e.g., caudate and putamen) support goal-directed action and the formation of stimulus-response habits (8) and are implicated in chronic and compulsive alcohol use in adults (9–12). These findings align with neurobiological models of addiction which propose that substance use progresses from reward-driven to habitual behavior, reflecting a ventral to dorsal striatal shift as use becomes more compulsive (13), yet this has not been investigated in a developmental context, limiting our ability to understand the emergence of alcohol use disorders.

Recent developmental neuroimaging suggests that corticostriatal circuits follow distinct maturational trajectories. Connectivity between decision making ventromedial prefrontal regions and the NAcc peaks during early adolescence, then decreases as control systems mature and risk taking decreases (14,15). In contrast, dorsal striatal connectivity tends to decrease gradually with age, reflecting increasing efficiency and specialization of action control (16). These developmental shifts are highly relevant to adolescent risk-taking, including alcohol use, as adolescence is marked by heightened reward sensitivity alongside still-maturing cognitive control systems (17), which may increase vulnerability to experimentation and early escalation (18). As reward-driven and habit-related systems have unique developmental trajectories (14–16), individual differences in these trajectories may shape when alcohol use begins and whether, and how rapidly, it escalates and remains elevated. However, most studies are either cross-sectional, have few measurement occasions, or focus on single regions or tasks, limiting insight into how these systems develop together over time. As a result, it remains unclear how normative trajectories of corticostriatal connectivity relate to key behavioral transitions, including the initiation of alcohol use and escalation in alcohol use frequency. Understanding these developmental processes is essential for linking neurobiological models of addiction to real-world substance use trajectories. Longitudinal approaches that examine corticostriatal connectivity may clarify how early reward-driven initiation evolves into habitual patterns of alcohol use and identify neural markers of risk during adolescence and early adulthood.

The present study addresses these gaps using data from the National Consortium on Alcohol and Neurodevelopment in Adolescence-Adulthood (NCANDA-A, https://ncanda.org/) cohort (19), a large longitudinal sample spanning adolescence through adulthood. We applied a data-driven, seed-based functional connectivity approach to characterize whole-brain corticostriatal connectivity with three striatal regions: the NAcc, caudate, and putamen. This approach enabled delineation of distinct developmental trajectories for each striatal region and examination of their relation to alcohol use behavior across adolescence and young adulthood. We hypothesized that NAcc connectivity would follow an inverted-U shaped trajectory, in line with literature showing heightened reward sensitivity during adolescence (20), whereas caudate and putamen connectivity would show gradual decreases with age, consistent with increasing efficiency of action control (16). We further predicted that alcohol use initiation would be associated with elevated NAcc connectivity, consistent with its role in reward-driven behavior, whereas stronger putamen connectivity would be associated with high alcohol use frequency and persistence through development. We also conducted substance-specificity analyses (binge drinking, nicotine, and cannabis) examining whether the observed brain-behavior associations generalized to other commonly used substances (reported in the Supplement). By integrating longitudinal neurodevelopmental trajectories with behavioral transitions in substance use, this study aims to clarify how corticostriatal maturation intersects with reward- and habit-related mechanisms of risk across adolescence and early adulthood. Understanding the developmental dissociation between ventral and dorsal striatal connectivity and how it relates to alcohol use can guide early identification of at-risk youth and inform interventions aimed at preventing the transition from experimental use to habitual drinking. This study represents one of the first longitudinal investigations of striatal network development in the context of initiation and maintenance of alcohol use.

## Methods

### Participants

NCANDA-A enrolled 831 participants (ages 12-21 at baseline) across five U.S. sites (Duke, Oregon Health & Science University, SRI International, University of California San Diego, and University of Pittsburgh) between 2013 and 2014. Participants had neuroimaging sessions annually until age 20 and biannually at 21 years and older. The present analyses included data from the first nine waves (baseline through Year 8 data release). Detailed recruitment and study procedures have been reported previously (19). Adult participants provided informed consent; minors provided assent with parental consent. All procedures were approved by the Institutional Review Board at each site. Nine participants were excluded due to lack of usable imaging data, yielding a final analytic sample of 822 participants and 4,358 scans. At baseline, 89% of participants met NIAAA criteria for no or low alcohol use.

### Measures

#### Alcohol use measures

Alcohol use was assessed annually using the Customary Drinking and Drug Use Record (21). Past-year alcohol use frequency was measured by the number of drinking days in the past year (0—365). Alcohol initiation was defined as the first assessment at which participants who reported no alcohol use at study entry subsequently reported ≥ 1 drinking day in the past year.

### Resting state connectivity

#### MRI acquisition and preprocessing

MRI data were collected across five NCANDA sites using 3T scanners. At UCSD, SRI, and Duke, imaging was performed on GE Discovery MR750 systems with 8-channel head coils; at OHSU and the University of Pittsburgh, data were acquired on Siemens Magnetom Tim Trio scanners with 12-channel coils and later on Siemens PRISMA FIT systems with 20-channel coils. Acquisition protocols were harmonized across sites and platforms and included high-resolution structural MRI, resting-state fMRI (TR = 2200 ms, TE = 30 ms; ∼10 min), and field maps for distortion correction. During resting-state scans, participants fixated on a gray screen with eyes open. Functional data were processed using a standardized pipeline designed to minimize motion and physiological artifacts (22). Volumes with framewise displacement > 0.3 mm were censored, and scans with > 30% censored volumes were excluded (145 observations). To account for site- and scanner-related variability, resting-state functional connectivity (RSFC) data were harmonized using neuroCombat (23).

#### Seed-based functional connectivity

Seed-based RSFC was computed for the nucleus accumbens (NAcc), caudate, and putamen, defined using the AAL3 anatomical atlas (24). For each seed, Pearson correlations were calculated between the mean seed time series and all brain voxels. Connectivity maps were then averaged within the Gordon 333-parcel cortical atlas (25), yielding one network per striatal seed. To characterize developmental trajectories, generalized additive mixed models (GAMMs) were fit separately for each parcel within each seeded network, with age modeled as a smooth term and random intercepts and slopes per participant. Statistical significance of age effects was assessed using Bonferroni correction within each seeded network. Trajectories were z-scored within seed to emphasize differences in developmental shape rather than absolute connectivity magnitude. Parcels exhibiting significant age-related change were clustered based on their age trajectories using k-means clustering. The clustering solution was evaluated using both the elbow method and silhouette criteria (26), with the goal of identifying a parsimonious and interpretable representation of the data; this procedure supported a two-cluster solution for each seed. Based on the k-means clustering results, FC values were computed by averaging connectivity across significant parcels within each cluster for each participant and time point, yielding six FC measures per observation (two clusters per striatal seed). All statistical analyses were conducted using R (v4.4.1; R Core Team 2024), and GAMMs were fit using the *mgcv* package (27). Code for the seed-based functional connectivity analyses can be found on Github (LabNeuroCogDevel/habit_ncanda).

### Statistical analysis

All statistical analyses used GAMMs to accommodate nonlinear developmental effects and repeated measures. Age was modeled using smooth terms to capture non-linear trajectories across adolescence and young adulthood. All models included random intercepts and slopes for participant and covaried for participant sex assigned at birth. Bonferroni correction was applied where noted. Formal specifications of all models can be found in the supplement. Code for all statistical analyses can be found on Github (LabNeuroCogDevel/habit_ncanda).

#### Alcohol use frequency trajectories

Developmental trajectories of alcohol use frequency were modeled using GAMMs with past-year alcohol use days as the outcome. Age was modeled as a smooth term, allowing us to capture nonlinear change over time. These models characterized normative patterns of alcohol use across development and provided context for subsequent initiation and maintenance analyses.

#### Association between functional connectivity and alcohol use initiation

Alcohol initiation was examined using a GAMM that included all participants. Initiation status was defined based on past-year alcohol use and coded as a three-level variable: pre-initiation (no reported alcohol use), post-initiation (any reported past-year alcohol use), and never initiated. GAMMs were fit with functional connectivity as the outcome and included a smooth term for age, and fixed effects of cluster membership, initiation status, and their interaction to test whether developmental connectivity trajectories differed before versus after initiation and relative to participants who never initiated alcohol use.

#### Association between functional connectivity and frequency of alcohol use

Frequency of alcohol use was examined using GAMMs fit as time-varying effect models (TVEMs) to capture age-dependent associations between functional connectivity and alcohol use frequency. For each cluster, alcohol use frequency (past-year drinking days) was modeled with a main smooth of age and a cluster-specific smooth interaction term, allowing the effect of functional connectivity on alcohol use vary across development. This approach enables identification of the developmental periods during which functional connectivity is most strongly associated with alcohol use. To account for multiple comparisons across clusters, Bonferroni correction was applied for each of the six clusters (two per seed).

#### Substance specificity analyses

To assess the substance specificity of observed associations, we conducted supplementary analyses applying the same analytic pipeline to other alcohol use behaviors (i.e., binge drinking) and other substance use outcomes (e.g., nicotine and cannabis); detailed measures and analytic specifications are provided in the Supplement.

## Results

### Participant demographics and alcohol use

Participant demographics at baseline are summarized in Supplementary Table 1. The final analytic sample included 822 participants (4,358 scans) ranging in age from 12.04 to 29.30 years at assessment (Figure 1a). Participants completed a mean of 5.30 study visits (median = 6). At the first available assessment, 32.50% of participants reported alcohol use (entered the study drinking). Among the remaining participants, 17.30% initiated alcohol use before age 18, 30.30% between ages 18 and 22, and 2.90% after age 22, while 17% never reported alcohol use at any assessment (Figure 1d). Among participants who initiated alcohol use during the study, the mean age of initiation was 18.9 years (SD = 2.07; range = 13.80–27.10 years; Figure 1c).

**Figure 1.**
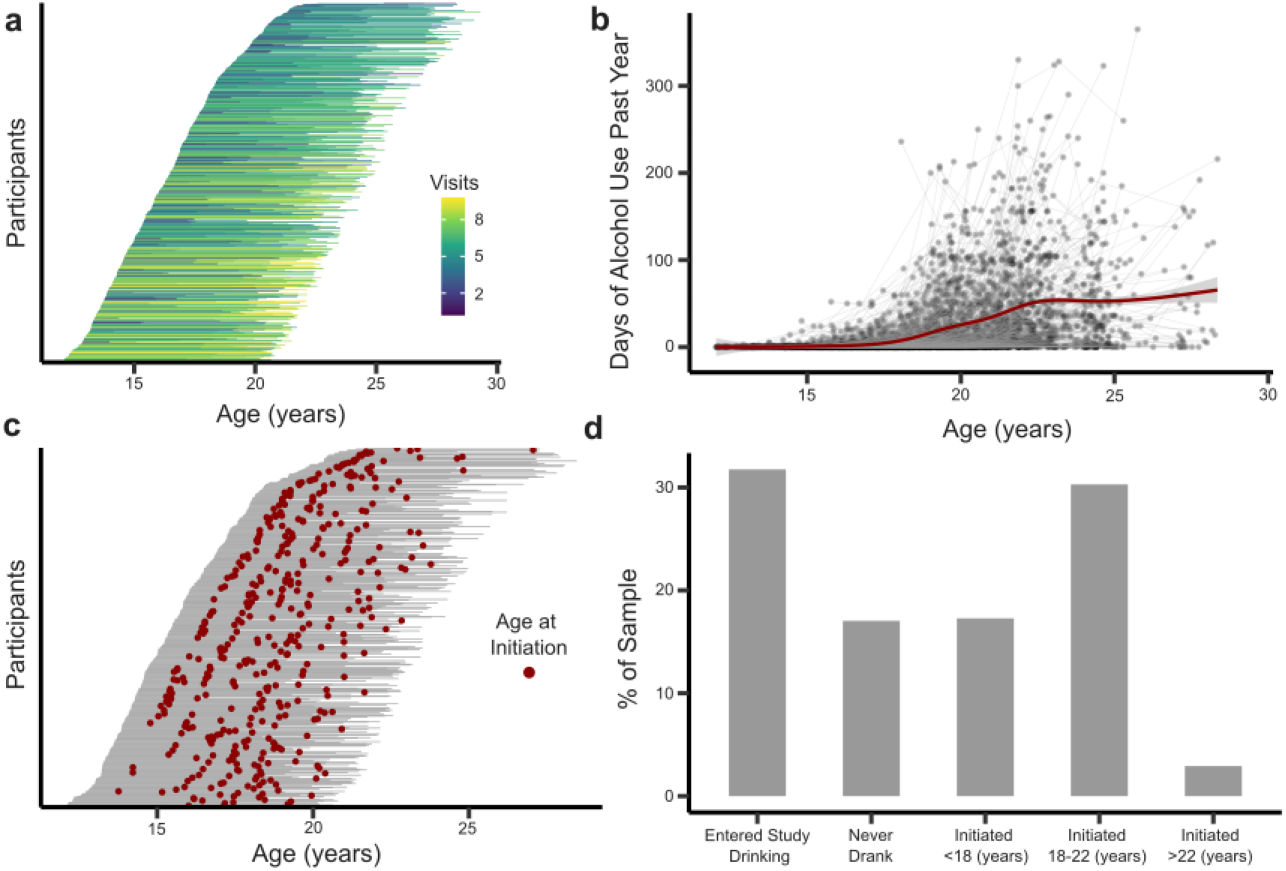
Sample characteristics and alcohol use patterns across development. **(a)** Age coverage and number of visits for the analytic sample. **(b)** Frequency of alcohol use across development. **(c)** Developmental timing of alcohol initiation. Red dots depict the visit where participants first self-reported alcohol use. **(d)** Sample composition by drinking status.

### Alcohol use frequency trajectories

Alcohol use frequency increased across adolescence and young adulthood, with trajectories varying across individuals (Figure 1b). The smooth term for age was significant and nonlinear (edf = 3.91, F = 423.1, p < 0.001), indicating a nonlinear increase in alcohol use over this developmental period. Inspection of the first derivative of the age smooth revealed a significant positive slope between ages 15.8 and 25.9 years, with the steepest increase at age 20.8, after which the slope remained positive but more gradual. No significant main effect of sex was observed. Random-effects estimates indicated substantial variability in individual trajectories, with between-subject differences in age-related change in drinking frequency, suggesting that participants differed not only in starting points but also in the rate of change across adolescence and young adulthood.

### Developmental trajectories of striatal seed-based functional connectivity

Seed-based functional connectivity analyses identified 142 clusters (56 from NAcc, 36 from caudate seed, and 50 from putamen seed) out of 999 that had significant developmental change after Bonferroni correction. Across striatal seeds, k-means clustering of age-related RSFC trajectories yielded two clusters per seed (Figure S1). For the NAcc, Cluster 1 comprised the majority of significant parcels (54 of 56), whereas Cluster 2 contained only two parcels. A similar pattern was observed for the caudate (30 vs. 6 parcels) and putamen (41 vs. 9 parcels).

Cluster membership showed clear correspondence with large-scale cortical networks (25). For the NAcc, clusters were dominated by parcels in the cingulo-opercular (n = 15) and auditory (n = 14) networks, with additional representation from sensorimotor (n = 7) and default mode regions (n = 5). Caudate clusters were primarily composed of cingulo-opercular parcels (n = 20), with comparatively limited representation from other networks (< 4 parcels each from other networks). For the putamen, clusters were again dominated by cingulo-opercular regions (n = 24), followed by auditory (n = 11) and sensorimotor (n = 11) networks (see Table S2).

Examination of the developmental trajectories revealed distinct age-related patterns across clusters. For clarity, we refer to each cluster by the characteristic shape of its trajectory: for the NAcc, the youth-peak cluster (formerly Cluster 1) and the decreasing cluster (formerly Cluster 2); for the caudate, the decreasing and increasing clusters (formerly Clusters 1 and 2, respectively); and for the putamen, the decreasing and inverted-sigmoidal clusters (formerly Clusters 1 and 2).

For the NAcc seed, the youth-peak cluster showed an inverted-U trajectory, increasing through mid-adolescence and declining in young adulthood, whereas the NAcc decreasing cluster exhibited a monotonic decrease across age (Figure 2a, 2d). For the caudate, the caudate decreasing cluster showed declining connectivity with age, while the caudate increasing cluster showed connectivity that rose from adolescence into adulthood (Figure 2b, 2e). For the putamen, both clusters exhibited decreasing connectivity, although the putamen inverted-sigmoidal cluster showed a trajectory that flattened in adulthood compared to the more uniformly declining putamen decreasing cluster (Figure 2c, 2f). Together, these patterns indicate that age-related changes in striatal connectivity vary by region and trajectory type, reflecting heterogeneous developmental pathways across cortical networks.

**Figure 2.**
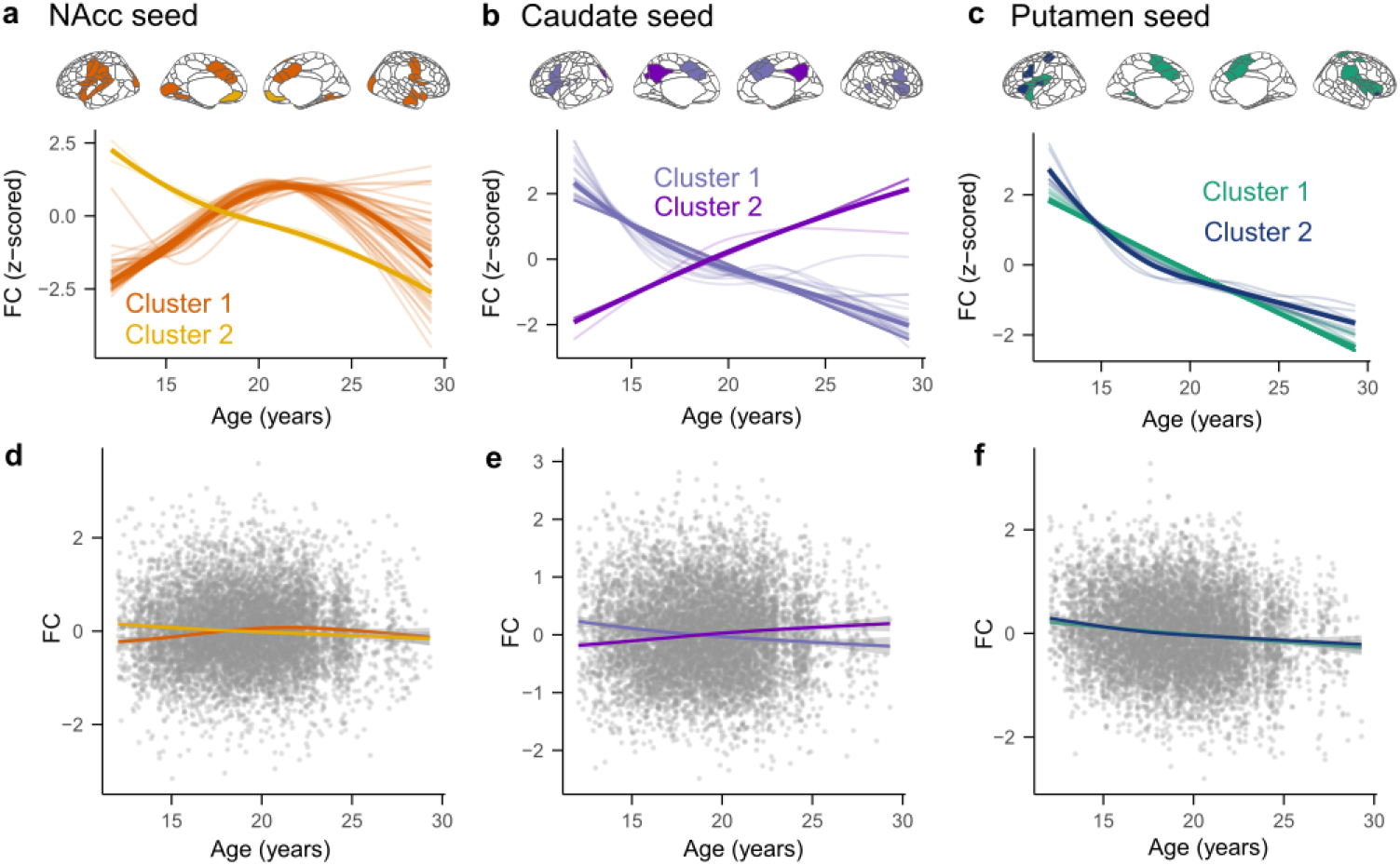
For all panels, parcels are colored according to the cluster to which they were assigned. Panels **(a–c)** display clustering results for each striatal seed (NAcc, caudate, putamen): the top row shows brain surface plots with parcels colored by cluster membership, and the bottom row shows developmental trajectories depicting the cluster centroids across age. Specifically, **(a)** NAcc clusters, **(b)** caudate clusters, and **(c)** putamen clusters. Panels **(d–f)** present the corresponding model-estimated trajectories overlaid on the raw data for each seed region (NAcc, caudate, putamen) to its respective cluster.

### NAcc and caudate connectivity is associated with alcohol use initiation

Developmental trajectories of striatal connectivity were further examined in relation to drinking initiation. GAMMs revealed significant main effects of drinking phase and seed-cluster, as well as significant interactions between seed-cluster and drinking phase (all *p* < 0.001). Post hoc comparisons using estimated marginal means indicated that for the NAcc youth peak cluster, connectivity was significantly higher after alcohol use relative to before initiation (Δ = -0.16, *p* < 0.001), suggesting an increase in NAcc connectivity in parallel with the onset of drinking (Figure 3a left). Similarly, connectivity in the caudate increasing cluster was significantly elevated after alcohol initiation compared to before (Δ = -0.14, *p* < 0.001; Figure 3b left). Together, these results indicate that both the NAcc youth-peak cluster and the caudate increasing cluster show heightened connectivity following alcohol initiation, highlighting a potential neural signature associated with early drinking. See Tables S3, S4, and Figure S2 for full results. No putamen clusters exhibited significant differences before/after initiation.

**Figure 3.**
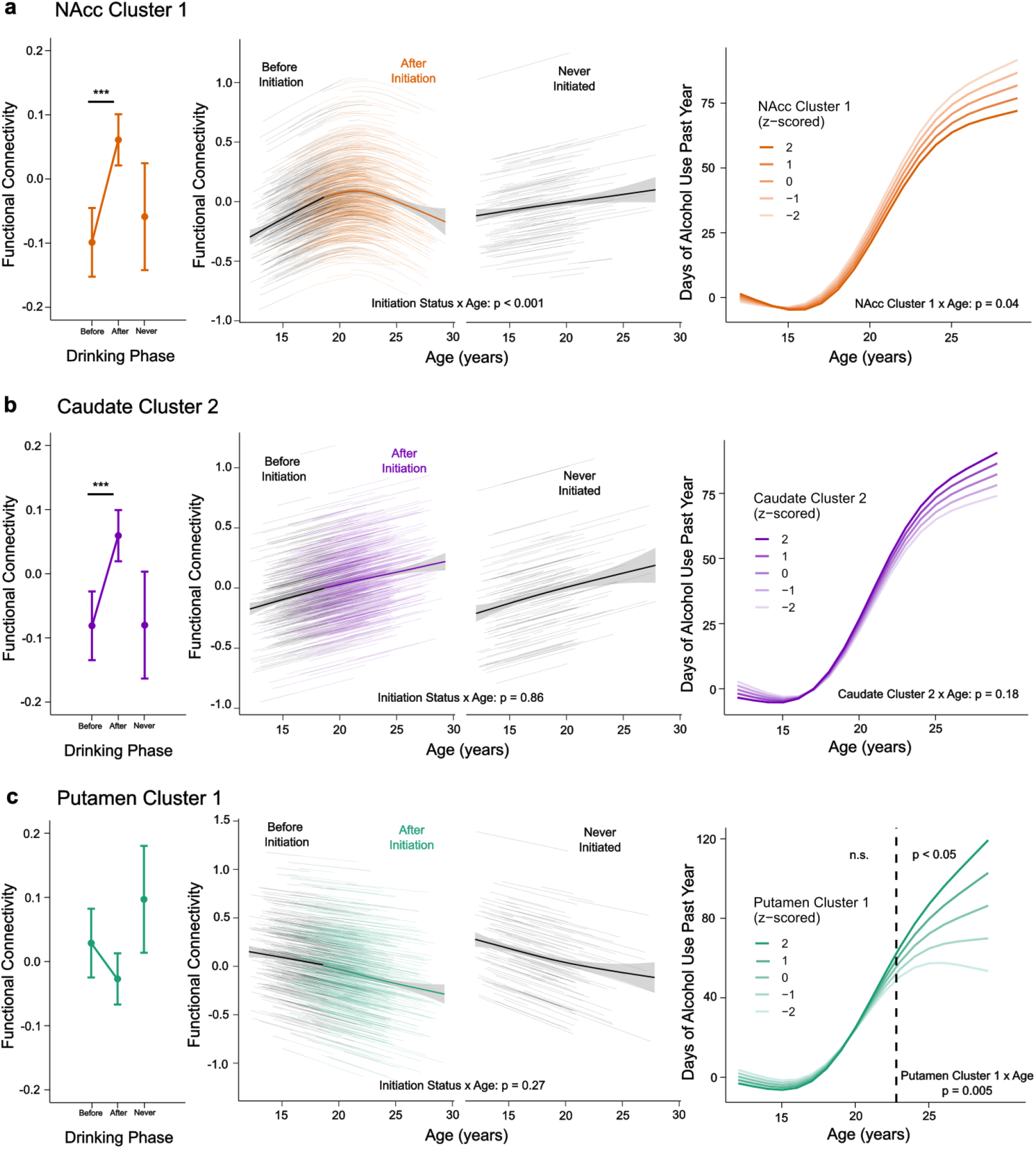
Relationship between corticostriatal connectivity and alcohol use initiation, drinking status, and frequency. *** = *p* < 0.001. **(a left)** Estimated marginal mean (± 95% CI) FC by alcohol initiation phase in NAcc Cluster 1. Horizontal brackets indicate pairwise comparisons among drinking-phase groups. **(a middle)** NAcc Cluster 1 FC trajectories based on whether an individual initiated alcohol use or if they never initiated alcohol use. Individual lines reflect model predictions for each participant, bold lines reflect the mean FC trajectory. **(a right)** NAcc Cluster 1 TVEM depicts associations between age and NAcc FC on days of alcohol use in the past year. **(b)** Caudate Cluster 2 results. Panels parallel those in (a). **(c)** Putamen Cluster 1 results. Panels parallel those in (a). The vertical dotted line (c right panel) represents the cutoff where the association between Putamen Cluster 1 FC and days of alcohol use in the past year is the strongest.

Following these significant effects, we examined whether developmental trajectories of striatal seed-based functional connectivity differed between individuals who self-reported alcohol use during the study and those who never reported alcohol use. To test this, we fitted two additional exploratory GAMMs for the NAcc youth peak cluster and caudate increasing cluster, specifying age smooths that differed by drinking group (self-reported drinking during the study vs. never drank). Drinking group was modeled as an ordered factor, such that the smooth term tested differences in age-related trajectories between groups. For the NAcc youth peak cluster, the parametric effect of drinking group was not significant; however, the difference between age smooths was significant (edf = 3.24, F = 10.78, p < 0.001), indicating that developmental changes in connectivity differed by drinking status. Visual inspection of the fitted trajectories showed that individuals who reported drinking during the study exhibited an inverted-U–shaped trajectory across age, whereas individuals who never reported drinking showed only a modest, monotonic increase in connectivity (Figure 3a center). These findings suggest that nonlinear developmental changes in NAcc connectivity were primarily driven by individuals who initiated alcohol use during the study. In contrast, for the caudate increasing cluster, there was no evidence that age-related trajectories differed by drinking group (edf = 1.00, F = 0.03, p = 0.86). Both groups showed similar age-related increases in connectivity, indicating no discernible moderation of caudate developmental trajectories by drinking status (Figure 3b center). See Figure S3 and Table S5 for full results.

### Putamen connectivity is associated with alcohol use frequency

To test whether striatal connectivity moderated age-related changes in alcohol use frequency, we fit a series of TVEMs using GAMMs, examining interactions between age and each of the six seed–cluster connectivity measures (Figure 3 right panels; Figure S4). After Bonferroni correction for six models (adjusted significance threshold *p* = 0.008), a significant age-by-connectivity interaction was observed for putamen decreasing cluster (edf = 3.37, F = 4.42, *p* = 0.005). Derivative-based inference of the interaction smooth indicated that higher connectivity in the putamen decreasing cluster was associated with a steeper age-related increase in past-year drinking beginning after approximately age 22 years (Figure 3c right). No other TVEM interactions survived correction for multiple comparisons. Full model results are provided in Supplementary Table S6. This finding suggests that age-related increases in putamen connectivity in young adulthood were associated with greater frequency of alcohol use across development.

### Binge drinking and substance specificity analyses

Specificity analyses revealed a consistent pattern across binge drinking, nicotine, and cannabis use. In each case, initiation of use was associated with increased functional connectivity in the NAcc youth peak cluster and the caudate increasing cluster. However, developmental trajectories of these circuits did not differ between individuals who initiated use and those who did not, and functional connectivity did not moderate age-related changes in use frequency for any substance (see Supplement).

## Discussion

This study examined the role of corticostriatal FC maturation in alcohol use initiation and trajectories of alcohol frequency from adolescence into adulthood. We first examined age-related changes in functional connectivity for each striatal seed (i.e., nucleus accumbens (NAcc), caudate, putamen, and cortical regions), identifying normative developmental patterns across the cortex. We then tested whether these developmental trajectories differed as a function of alcohol use initiation and whether connectivity strength was related to use frequency. We show that connectivity patterns varied systematically across striatal subdivisions and differentially contributed to substance use patterns: NAcc connectivity followed an inverted U-shaped trajectory, peaking during adolescence; caudate connectivity showed a monotonic decrease with age; and putamen connectivity showed a similar monotonic decreasing pattern. Importantly, after controlling for age, NAcc connectivity exhibited a post-initiation increase, while in adulthood, higher putamen connectivity was linked to more frequent alcohol use. To our knowledge, no prior longitudinal study has simultaneously examined both reward-driven and habitual neural circuit function as predictors of alcohol use initiation and subsequent frequency across adolescence and into adulthood.

### Maturation of corticostriatal functional connectivity from adolescence to adulthood

The corticostriatal clusters identified in the present study primarily exhibited functional connectivity with regions of the cingulo-opercular network, recently reframed as the action-mode network (AMN), a system implicated in sustained cognitive control, performance monitoring, and the maintenance of goal-directed behavior (28). Our findings suggest that developmental changes in striatal connectivity may reflect shifting interactions between motivational and action-control systems across adolescence and early adulthood. Specifically, connectivity between the NAcc and parcels in the AMN followed an inverted U-shaped trajectory, peaking in late adolescence before declining into adulthood. This pattern has been found in previous studies (15,29) and is broadly consistent with developmental models proposing heightened reward salience and motivational drive during adolescence (30), followed by increasing regulatory stability with maturation. In contrast, connectivity between the putamen and action-mode regions showed a monotonic decrease with age, which has also been found in past studies (29,31). Elevated putamen coupling earlier in adolescence may reflect greater engagement of monitoring and top-down control processes as goal-directed and stimulus–response behaviors are acquired. Subsequent reductions in connectivity may indicate increasing efficiency and autonomy of putamen circuits as behaviors become more automatized and require less sustained control. Together, these dissociable developmental trajectories support neurobiological models in which ventral and dorsal striatal systems mature along distinct timelines (32), potentially contributing to the transition from reward-guided exploration toward more habitual behavioral control.

### Increased NAcc functional connectivity relates to alcohol use initiation

Alcohol use initiation was associated with elevated functional connectivity between the NAcc and regions of the AMN, consistent with past research (6,7,33). These post-initiation increases in AMN-NAcc connectivity may reflect a strengthening of the coupling between reward signals and sustained control processes, potentially enhancing the motivational salience of alcohol-related cues (34) in the context of elevated risk-taking (35), or supporting early reinforcement-related consolidation of alcohol-guided action strategies (13,36). Such increases may also bias behavior toward exploratory or novelty-seeking actions, particularly among youth with higher baseline risk-taking tendencies (35), thereby facilitating subsequent experimentation. By highlighting changes in large-scale network integration following initiation, the present results extend prior work focused primarily on reward responsivity by suggesting that altered coordination between reward and control systems may play a key role in the transition from initial use to more goal-directed alcohol engagement. Elevated AMN-NAcc connectivity in this context may represent a reinforcement-related shift in coupling that emerges after drinking begins. From a developmental perspective, this pattern may represent a neural marker of vulnerability during adolescence when motivational systems are highly active but regulatory control is still maturing (17). Identifying such network-level differences is important because early initiation is a robust predictor of later alcohol-related problems (4), and alterations in reward–control circuitry may represent a target for early prevention, intervention, and treatment efforts (37–39).

### Developmental trajectories of NAcc FC differed as a function of alcohol use

An additional finding was that youth who did not initiate alcohol use showed a gradual increase in NAcc connectivity across adolescence and early adulthood. Given that alcohol experimentation is common during adolescence (2), the inverted U-shaped trajectory may reflect a more typical developmental pattern of reward–control integration during this period. The peak in connectivity during adolescence could correspond to heightened coordination between motivational and action-control systems, followed by refinement and increased efficiency as maturation progresses. In contrast, the continued linear increase observed among non-initiators may reflect delayed experiential engagement with reward-related behaviors or alternative later developmental timing of corticostriatal maturation. Importantly, these findings do not imply that alcohol use implicitly causes functional neural reorganization; rather, they suggest that differences in behavioral experience may coincide with distinct trajectories of reward-related network development. Such heterogeneity underscores the importance of considering developmental timing and common adolescent experiences when interpreting neural correlates of alcohol use.

### Putamen functional connectivity relates to the frequency of alcohol use

The present findings are broadly consistent with theoretical models of addiction emphasizing drug-induced neuroadaptations in corticostriatal circuitry that shift behavioral control from reward-driven to habitual processing (40,41). Within this framework, habit formation has been proposed to play a central role in the progression of substance use through stronger dorsal striatal circuits that support automatic, cue-driven behavior. Consistent with this account, we found that more regular alcohol use was associated with attenuated age-related declines in putamen FC, such that individuals with more frequent drinking exhibited relatively elevated connectivity in young adulthood. This attenuated decrease in connectivity may reflect differences in the maturation of neural systems supporting habitual behavior. One possibility is that elevated connectivity in this circuit reflects greater engagement of habit-related processes that promote more automatic, stimulus–response alcohol use. Alternatively, this pattern may indicate a less mature or more reward-driven control system, whereby individuals continue to recruit dorsal striatal circuitry to regulate behavior in the presence of alcohol-related cues. The heightened putamen connectivity observed here may therefore reflect increased reliance on reward systems rather than fully consolidated habitual control. These findings suggest that individual differences in alcohol use during young adulthood may be associated with altered developmental trajectories in habit-related circuitry, potentially marking a period of increased vulnerability for the consolidation of more automatic and persistent patterns of alcohol use.

Time-varying effect models revealed that the association between putamen FC and drinking days was strongest after approximately age 22, a developmental period corresponding to major life transitions such as completing college and entering the workforce, when drinking often shifts from social to more regular or solitary use (42). Elevated connectivity in habit-related circuitry during this window may reflect increased consolidation of alcohol use patterns, potentially marking a neurobiological inflection point in the transition from moderate to heavier and sustained drinking. These findings align with prior work showing greater dorsal striatal activation in heavy versus light drinkers, supporting a shift from ventral to dorsal striatal control as alcohol use escalates (43), and with studies reporting increased reliance on stimulus-response habit learning in adults with alcohol use disorder (AUD), accompanied by reduced engagement of goal-directed circuitry and heightened putamen recruitment (44). Converging neuroimaging findings further demonstrate hyperconnectivity between the putamen and cortical regions as alcohol use severity increases in adulthood (45,46). The present longitudinal results extend this literature by showing that functional connectivity within habit-related circuits differentiates drinking patterns in young adults and varies across development.

### Binge drinking and substance specificity analyses

To clarify the specificity of the observed brain–behavior associations, we examined whether the corticostriatal connectivity effects identified for alcohol frequency generalized to binge drinking, nicotine, and cannabis use. These substance-specificity analyses revealed a consistent pattern at the stage of initiation: across binge drinking, nicotine, and cannabis, first use was associated with increased functional connectivity within striatal circuits involving the NAcc and caudate. This convergence suggests that initiation-related connectivity changes may reflect a transdiagnostic developmental or behavioral transition, potentially linked to heightened risk taking, novelty seeking, or sensitivity to salient rewards, that engages corticostriatal systems similarly across substances.

In contrast, associations between striatal connectivity and frequency or persistence of use were observed only for alcohol use (but not binge drinking episodes), suggesting that certain corticostriatal connectivity features may show substance-specific associations with sustained use. This dissociation supports a framework in which initiation effects reflect shared NAcc mediated developmental mechanisms, whereas neural signatures of continued use may depend more strongly on substance-specific factors such as pharmacology, dosing patterns, and neuroadaptations. Notably, it is possible that alcohol exerts particularly pronounced effects on corticostriatal circuitry during the transition from experimental to regular use, highlighting a potential window of heightened neurobiological vulnerability.

### Limitations

There are several limitations worth noting. First, the approximately one-year spacing between study visits (and two years between neuroimaging visits in 21-year-old participants and older) makes it difficult to determine the precise timing of alcohol initiation relative to measured brain connectivity. As a result, we cannot infer whether the observed FC differences represent premorbid risk markers or reflect early neurobiological changes following initial alcohol use. Future work would benefit from collecting data at shorter intervals or incorporating ecological momentary assessment (EMA). EMA can capture substance-use behavior in real time or near real time, reducing recall bias, improving temporal precision, and allowing researchers to link fluctuations in daily experiences, mood, and context with substance-use events (47). Integrating EMA closer to imaging sessions could substantially increase the temporal specificity needed to identify when alcohol use escalates and how it relates to dynamic neural changes. Second, we did not examine whether the FC patterns identified here predict longer-term outcomes, such as subsequent escalation trajectories or later alcohol use disorder (AUD). Future longitudinal studies should evaluate whether connectivity profiles in NAcc-, caudate-, or putamen-based clusters prospectively predict AUD diagnoses or related clinical impairments. Third, the group of individuals who abstain from alcohol is likely heterogeneous. Reasons for abstaining, including familial risk of problematic use, cultural or religious norms, health concerns, or personal temperament, may differ substantially across participants, and such heterogeneity could obscure meaningful brain–behavior patterns or introduce confounding factors unrelated to alcohol exposure itself. Additionally, our longitudinal coverage of abstainers is limited beyond age 20, leaving open the question of whether a more fully powered sample of individuals with zero alcohol exposure would show similar late-adolescent decreases in functional connectivity to those observed among drinkers toward the upper end of the study period. As a result, it remains unclear whether the divergence in connectivity patterns reflects alcohol-specific effects or broader developmental trajectories that we were underpowered to detect within the abstaining group. Finally, the observational design also limits causal inference, as unmeasured environmental, genetic, or psychosocial factors may influence both FC development and drinking behaviors.

### Conclusion

In summary, the current study found that the maturation of distinct corticostriatal circuits supports different stages of alcohol use. Reward-related connectivity in the NAcc following adolescent initiation may reflect processes associated with early experimentation due to elevated sensation seeking during the developmental period, whereas elevated putamen connectivity in young adulthood may support the emergence of more habitual or frequent patterns of use. These results highlight how ventral and dorsal striatal circuits follow unique developmental trajectories that interact with behavioral transitions to shape substance use risk. Finally, these findings underscore the importance of longitudinal designs that span adolescence to adulthood for characterizing brain–behavior associations in substance use and identifying markers that predict more problematic patterns of substance use into adulthood. By integrating longitudinal developmental change with alcohol use patterns, this study provides a framework for understanding the neurobiological mechanisms underlying the transition from alcohol use experimentation to more regular use patterns, with potential implications for interventions targeting critical developmental windows before patterns of alcohol use become more entrenched.

