## Supplement for "Neurodevelopment of Corticostriatal Circuits and Risk for Alcohol Use During the Transition from Adolescence to Adulthood"

**Supplement for Petrie et al., *Neurodevelopment of Corticostriatal Circuits and Risk for Alcohol Use During the Transition from Adolescence to Adulthood* for submission to the *American Journal of Psychiatry***

**Supplemental Methods**

**Participants**

The National Consortium on Alcohol and NeuroDevelopment in Adolescence-Adulthood (NCANDA-A) is an ongoing accelerated longitudinal study of healthy 12-22 year olds (1), that examines how individual difference in neurodevelopment are associated with alcohol use problems, and how alcohol use effects the brain across development. NCANA-A enrolled 831 participants (ages 12-21 baseline) across five U.S. sites (Duke, Oregon Health & Science University, SRI International, University of California San Diego, and University of Pittsburgh). Each site recruited participants to reflect the racial and ethnic composition of its local community. NCANDA-A recruited youth with minimal prior substance exposure. At the same time, the sampling strategy enriched the cohort for individuals at elevated risk for alcohol use disorder (AUD), such as those with a family history of substance use disorder, (SUD) resulting in approximately 47% of the sample being classified as higher risk. Exclusion criteria included contraindications to MRI (e.g., claustrophobia, pregnancy, or non-removable metal), medical conditions that could affect neuroimaging (e.g., head injury with loss of consciousness), current or persistent psychiatric disorders likely to interfere with study participation (e.g., psychotic disorders), and the use of psychiatric medications (1). Participant demographic information is provided in Table S1.

**MRI acquisition and preprocessing**

Detailed methods for MRI acquisition have been reported elsewhere (2,3). High-resolution T1-weighted structural images were acquired across five sites using 3T MRI systems from two manufacturers. Three sites used GE Healthcare Discovery MR750 scanners (University of California San Diego, SRI International, and Duke University), and two sites used Siemens Healthineers TIM TRIO systems (University of Pittsburgh and Oregon Health and Science University). GE sites employed an inversion recovery spoiled gradient recalled (IR-SPGR) sequence with parallel imaging (TR, 6 ms; TE, 2 ms; flip angle, 11°; matrix, 256 x 256; FOV, 24cm; 1.2 x .94 x .94 mm voxel size, 146 slices), whereas Siemens sites acquired T1-weighted images using a magnetization-prepared rapid gradient echo (MPRAGE) sequence (TR, 6 ms; TE, 3 ms; flip angle, 9°; matrix, 256 x 256; FOV, 24 cm; 1.2 x .94 x .94 mm voxel size, 160 slices). Acquisition parameters were harmonized across sites to ensure comparable spatial resolution (approximately 1 mm in-plane). Scanner stability was monitored using daily phantom scans.

Structural images were preprocessed to remove non-brain tissue and were normalized to standard Montreal Neurological Institute (MNI) space using both linear (FLIRT) and nonlinear (FNIRT) registration. Resting-state functional MRI data were processed using a standardized pipeline designed to minimize motion and physiological artifacts (Hallquist et al., 2013). Preprocessing steps included slice-timing and motion correction, skull stripping, intensity normalization, despiking, and spatial smoothing (5 mm Gaussian kernel). Functional images were then coregistered to the structural image and warped to MNI space. Nuisance regression included head motion parameters and their derivatives, as well as white matter and cerebrospinal fluid signals. Bandpass filtering (0.009–0.08 Hz) was applied concurrently with nuisance regression.

**K-means trajectory clustering methods**

To identify spatially distributed patterns of developmental change in seed-based functional connectivity, we applied k-means clustering to model-derived age trajectories. For each seed region, generalized additive mixed models (GAMMs) were first used to estimate age-related trajectories of connectivity between the seed and all cortical regions defined by the Gordon Atlas (4). To focus clustering on developmentally dynamic connections, Bonferroni correction was applied across models for each seed. This resulted in 142 of 999 total seed–target connections (333 parcels per seed) showing significant age-related change, including 56 parcels for the nucleus accumbens, 50 for the putamen, and 36 for the caudate. Predicted connectivity values were then extracted across a uniform age grid for each significant seed–target pair, yielding smooth trajectories that characterized age-related changes in functional connectivity.

These predicted trajectories were then reorganized into a matrix in which each row represented a seed–target trajectory and each column corresponded to an age point. To ensure that clustering reflected differences in trajectory shape rather than absolute connectivity magnitude, each trajectory was standardized (z-scored) across age. This procedure emphasized relative developmental patterns and minimized the influence of baseline differences in connectivity strength.

K-means clustering was applied separately for each seed region to group trajectories with similar developmental profiles. The algorithm was implemented using multiple random initializations (nstart = 25) to reduce sensitivity to starting values and to ensure convergence on stable solutions. The optimal number of clusters (k) was determined using a combination of theoretical considerations, visualization of developmental trajectories, and quantitative criteria including elbow plots of within-cluster sum of squares and silhouette analysis. Final cluster solutions were selected based on convergence across these approaches and interpretability of developmental patterns.

Cluster assignments were subsequently mapped back to the original connectivity data. For each participant at each time point, functional connectivity values were labeled according to the cluster membership of the corresponding seed-parcel trajectory. This procedure enabled downstream statistical modeling of connectivity development at the level of cluster-defined networks rather than individual regions, thereby reducing dimensionality while preserving developmental heterogeneity.

**Statistical analysis**

All statistical analyses were conducted using R Statistical Software (v4.5.0; R Core Team 2025). Generalized Additive Mixed Models (GAMMs) were used to capture any non-linear effects and account for repeated measures. All GAMM models were fit using the *mgcv* package (5).

***Alcohol use frequency trajectories***

We characterized alcohol use trajectories using a GAMM, and were specified as follows:

${Alcohol Use Freq}_{i,j}= \beta_{0}+ \beta_{1}\left( {sex}_{i} \right)+ f_{1}\left( {age}_{i,j} \right)+ b_{0i}+ b_{1i}{age}_{i,j}+ \varepsilon_{i,j}$	(1)

where ${Alcohol Use Freq}_{i,j}$ represents the number of self-reported alcohol use days in the past year for visit *j* and person *i*; $\beta_{0}$ is the fixed intercept; $\beta_{1}\left( {sex}_{i} \right)$ is a fixed effect for sex assigned at birth; $f_{1}\left( {age}_{i,j} \right)$ is a penalized smooth function for age with the maximum knots set to 5; $b_{0i}$ is the random intercept term for each participant; $b_{1i}{age}_{i,j}$ is the random slope for age; and $\varepsilon_{i,j}$ is the residual error term. All random effects and residuals are assumed to be independent and normally distributed with a mean of 0 and constant variance.

***Association between functional connectivity and alcohol use initiation***

We then examined how each cluster (FC Cluster; Six clusters across the three seeds, two clusters per seed) and alcohol use phase (e.g., before initiation, after initiation, and never initiated) were associated with changes in functional connectivity. The model was specified as follows:

${FC}_{i,j,k}= \beta_{0}+ \beta_{1}\left( {sex}_{i} \right)+ \beta_{2}\left( {FC Cluster}_{k} \right)+ \beta_{3}\left( {Phase}_{i,j} \right)+ \beta_{4}\left( {{FC Cluster}_{k} \times Phase}_{i,j} \right)+ f_{1}\left( {age}_{i,j} \right) +b_{0i}+ b_{1i}{age}_{i,j}+ \varepsilon_{i,j,k}$	(2)

where ${FC}_{i,j,k}$ represents functional connectivity for FC Cluster *k* at visit *j* for person *i*; $\beta_{2}\left( {FC Cluster}_{k} \right)$ is the fixed effect for each FC Cluster; $\beta_{3}\left( {Phase}_{i,j} \right)$ is the fixed effect for each drinking phase; and $\beta_{4}\left( {{FC Cluster}_{k} \times Phase}_{i,j} \right)$ is the fixed effect interaction between FC Cluster and drinking phase. All other variables are specified identically as Equation 1.

***Developmental trajectories of functional connectivity based on alcohol use***

Following these significant effects, we examined whether developmental trajectories of striatal seed-based functional connectivity differed between individuals who self-reported alcohol use during the study and those who never reported alcohol use. Age-related effects were modeled using smooth functions, and time-varying differences between drinking groups were assessed using ordered-factor smooth interactions. The model was specified as follows:

${FC}_{i,j}= \beta_{0}+ \beta_{1}\left( {sex}_{i} \right)+ \beta_{2}G_{i}+ f_{1}\left( {age}_{i,j} \right) +f_{1}\left( {age}_{i,j} \right) \times G_{i}+ b_{0i}+ b_{1i}{age}_{i,j}+ \varepsilon_{i,j,k}$	(3)

where $\beta_{2}G_{i}$ represents the fixed effect for drinking group; $f_{1}\left( {age}_{i,j} \right)$ represents the reference smooth trajectory for the never initiated alcohol use group; and $f_{1}\left( {age}_{i,j} \right)$ represents the difference in the smooth trajectory for the initiated group. All other variables are specified identically as Equation 1. For the initiated group, the trajectory parameter can be described as:

$f_{Initiated}\left( {age}_{i,j} \right)= f_{Never}\left( {age}_{i,j} \right)+ f_{Diff}\left( {age}_{i,j} \right)$	(4)

where the initiated group trajectory can be described as a sum of the never initiated trajectory and the difference trajectory.

***Association between functional connectivity and frequency of alcohol use***

We were also interested in whether FC moderated the association between age and frequency of alcohol use. We used a time-varying effect model (TVEM) to examine how the association between striatal functional connectivity and alcohol use frequency change across age. TVEM allows the effect of a predictor to vary smoothly over continuous time, rather than assuming a fixed effect. This allowed us to examine the age range where the association between seeded connectivity and alcohol use frequency was the strongest. The model was specified as follows:

${Alcohol Use Freq}_{i,j}= \beta_{0}+ \beta_{1}\left( {sex}_{i} \right)+ f_{1}\left( {age}_{i,j} \right)+ f_{2}\left( {age}_{i,j} \right) \times{FC cluster}_{i,j}+ b_{0i}+ b_{1i}{age}_{i,j}+ \varepsilon_{i,j}$	(5)

where $f_{2}\left( {age}_{i,j} \right) \times{FC cluster}_{i,j}$ represents the time-varying effect of FC for each cluster. If significant, we then calculated the first derivative of the TVEM effect of functional connectivity on alcohol use to examine how this association changed across development.

**Supplemental Results**

**K-means clustering results**

To characterize heterogeneity in developmental functional connectivity (FC) patterns, we applied k-means clustering separately to longitudinal FC estimates derived from each striatal seed (NAcc, caudate, and putamen).

For the nucleus accumbens (NAcc), a two-cluster solution was selected based on elbow and silhouette metrics, as well as visual inspection of developmental trajectories. The resulting clusters reflected distinct age-related patterns of functional connectivity. Visualization in two-dimensional space demonstrated clear separation between clusters, supporting the stability of the solution (Figure S1a). Cluster 1 exhibited a nonlinear, inverted U-shaped trajectory across development, whereas Cluster 2 showed a monotonic decrease in connectivity with age. For the caudate, clustering also identified two groups with separable FC trajectories. A two-cluster solution was selected based on elbow and silhouette metrics, and visual inspection of developmental trajectories. Cluster 1 exhibited a monotonic decrease in connectivity with age, and Cluster 2 exhibited a monotonic increase in connectivity with age (Figure S1b). For the putamen, a two-cluster solution was also selected. Although modest improvements in within-cluster Sum of Squares and silhouette width were observed with additional clusters, these gains were minimal and did not correspond to meaningful qualitative differences in trajectory shape. Inspection of the trajectories revealed largely similar monotonic decreases in connectivity with age across higher cluster solutions. Therefore, a two-cluster solution was retained to balance interpretability and parsimony (Figure S1c).

**Substance Specificity Analyses**

To assess the substance specificity of observed brain–behavior associations, we applied the same analytic pipeline to other substance‑use outcomes. We used data from the Customary Drinking and Drug Use Record for all specificity analyses. Binge drinking frequency was measured by the number of self-reported binge drinking episodes in the past year (4+ drinkers within an occasion for females, and 5+ drinks within an occasion for males). Binge drinking initiation was defined as the first assessment at which participants who reported no binge drinking at study entry subsequently reported ≥ 1 binge drinking episode in the past year. For nicotine frequency and initiation, we used monthly retrospective self‑report of past‑month use. We did not include past‑year nicotine measures because the wording of the item was ambiguous, potentially referring either to the number of cigarettes smoked in the past year or to the number of days of use, limiting interpretability. Past‑month nicotine use provided a clearer and more reliable measure. For cannabis frequency and initiation, we used retrospective self‑report of past‑month and past‑year use.

***Binge Drinking Specificity Analyses***

Developmental trajectories of striatal connectivity were further examined in relation to binge drinking initiation. GAMMs revealed significant main effects of drinking phase and seed-cluster, as well as significant interactions between seed-cluster and drinking phase (all *p* < 0.001). Post hoc comparisons using estimated marginal means indicated that for the NAcc youth peak cluster, connectivity was significantly higher after alcohol use relative to before initiation (Δ = -0.12, *p* < 0.001), suggesting an increase in NAcc connectivity in parallel with the onset of drinking (Figure S5a left). Similarly, connectivity in the caudate increasing cluster was significantly elevated after alcohol initiation compared to before (Δ = -0.14, *p* < 0.001; Figure S5a left).

Following these significant effects, we examined whether developmental trajectories of striatal seed-based FC differed between individuals who self-reported binge drinking during the study and those who never reported binge drinking. To test this, we fitted two additional exploratory GAMMs for the NAcc youth peak cluster and caudate increasing cluster, specifying age smooths that differed by binge drinking group. For both NAcc youth peak cluster and caudate increasing cluster, there was no evidence that age related trajectories differed by binge drinking group (both *p’s* > 0.05; Figure S5 right). Both groups showed similar age-related increases in connectivity, indicating no discernable moderation of NAcc or caudate developmental trajectories by binge drinking status.

To test whether striatal connectivity moderated age‑related changes in binge drinking frequency, we fit a series of time‑varying effect models (TVEMs) examining interactions between age and each of the six seed‑cluster connectivity measures. No TVEM interactions were significant and are therefore not discussed further.

***Nicotine Specificity Analyses***

Developmental trajectories of striatal connectivity were also examined in relation to nicotine use initiation based on past‑month self‑report. GAMMs revealed significant main effects of nicotine use phase and seed‑cluster, as well as significant interactions between seed‑cluster and nicotine use phase (all p < 0.001). Post hoc comparisons using estimated marginal means indicated that for the NAcc youth peak cluster, connectivity was significantly higher after nicotine initiation relative to before (Δ = -0.21, p < 0.001; Figure S6a left). Similarly, connectivity in the caudate increasing cluster was significantly elevated following nicotine initiation compared to pre‑initiation levels (Δ = -0.16, p < 0.001; Figure S6b left).

Following these effects, we examined whether developmental trajectories of striatal seed‑based FC differed between individuals who reported any past‑month nicotine use during the study and those who never reported nicotine use. Exploratory GAMMs specifying age smooths that differed by nicotine use group provided no evidence that age‑related trajectories differed between groups for either the NAcc youth peak cluster or the caudate increasing cluster (both p’s > 0.05). Both groups showed comparable age‑related increases in connectivity, indicating no moderation of striatal developmental trajectories by nicotine use status (Figure S6 right).

To test whether striatal connectivity moderated age‑related changes in nicotine use frequency, we fit a series of time‑varying effect models (TVEMs) examining interactions between age and each of the six seed‑cluster connectivity measures. No TVEM interactions were significant and are therefore not discussed further.

***Cannabis Specificity Analyses***

We next examined developmental trajectories of striatal connectivity in relation to cannabis use initiation using both past‑month and past‑year self‑report measures. GAMMs revealed significant main effects of cannabis use phase and seed‑cluster, as well as significant interactions between seed‑cluster and cannabis use phase (all p < 0.001). Post hoc estimated marginal means indicated that connectivity within the NAcc youth peak cluster was significantly higher following cannabis initiation relative to pre‑initiation (Past month: Δ = -0.14, p < 0.001, Figure S7a left; Past year: Δ = -0.14, p < 0.001, Figure S8a left). Similarly, connectivity in the caudate increasing cluster was significantly elevated after cannabis initiation compared to before (Past month: Δ = -0.14, p < 0.001, Figure S7b left; Past year: Δ = -0.12, p < 0.001, Figure S8b left).

As with alcohol and nicotine, we tested whether age‑related trajectories of striatal seed‑based FC differed between individuals who reported cannabis use during the study and those who never reported cannabis use. Exploratory GAMMs with group‑specific age smooths provided no evidence that developmental trajectories differed by cannabis use status for either the NAcc youth peak cluster or the caudate increasing cluster (both p’s > 0.05). Both groups demonstrated similar age‑related increases in connectivity (Figure S7 right, Figure S8 right).

Finally, we tested whether striatal connectivity moderated age‑related changes in cannabis use frequency using TVEMs examining age × connectivity interactions across the six seed‑cluster measures. No significant interactions were observed for either past‑month or past‑year cannabis use, and these results are therefore not discussed further.

**Supplemental Tables**

**Table S1.** Demographic information at baseline for the current study can be found in Table S1.

|  | **N** | **%** |
| --- | --- | --- |
| **Sex assigned at birth** |  |  |
| Female | 417 | 51% |
| Male | 405 | 49% |
| **Race/Ethnicity** |  |  |
| Asian | 60 | 7% |
| Hispanic | 97 | 12% |
| Non-Hispanic Black/African American | 96 | 12% |
| Non-Hispanic White | 527 | 64% |
| Native American/Pacific Islander/Other | 42 | 5% |
| **Highest household parent income** |  |  |
| < 50k | 126 | 15% |
| 50k – 100k | 196 | 24% |
| > 100k | 414 | 50% |
| Don’t Know | 22 | 3% |
| **Highest household parent education** |  |  |
| High school Diploma/GED/Associate Degree | 125 | 15% |
| Bachelor’s Degree | 259 | 32% |
| Post-Graduate Degree | 355 | 43% |
| Other/None of the above | 23 | 3% |

**Table S2.** Significant parcels after Bonferroni correction by seed, network, and cluster assignment

| **Seed** | **Cluster Assignment** | **Functional Network** | **# of Parcels** |
| --- | --- | --- | --- |
| NAcc | 1 | Auditory | 14 |
|  |  | Cingulo-Opercular | 15 |
|  |  | Default | 3 |
|  |  | None | 5 |
|  |  | Salience | 2 |
|  |  | Sensorimotor Hand | 1 |
|  |  | Sensorimotor Mouth | 6 |
|  |  | Ventral Attention | 1 |
|  |  | Visual | 7 |
|  | 2 | Default | 2 |
| Caudate | 1 | Auditory | 3 |
|  |  | Cingulo-Opercular | 20 |
|  |  | Dorsal Attention | 3 |
|  |  | Frontal Parietal | 1 |
|  |  | Salience | 2 |
|  |  | Ventral Attention | 1 |
|  | 2 | Default | 3 |
|  |  | Medial Parietal | 1 |
|  |  | None | 1 |
|  |  | Visual | 1 |
| Putamen | 1 | Auditory | 9 |
|  |  | Cingulo-Opercular | 23 |
|  |  | Sensorimotor Hand | 6 |
|  |  | Sensorimotor Mouth | 2 |
|  |  | Visual | 1 |
|  | 2 | Auditory | 2 |
|  |  | Cingulo-Opercular | 1 |
|  |  | Dorsal Attention | 2 |
|  |  | Salience | 1 |
|  |  | Sensorimotor Hand | 2 |
|  |  | Sensorimotor Mouth | 1 |

**Table S3.** Association between seed-based functional connectivity and alcohol use initiation.

| **Parametric coefficients** |  |  |  |
| --- | --- | --- | --- |
|  | **Estimate** | **S.E.** | ***p*** |
| Intercept | -0.17 | 0.03 | < 0.001*** |
| Sex (Male) | 0.16 | 0.03 | < 0.001*** |
| Nacc Cluster 2 | 0.10 | 0.03 | < 0.001*** |
| Caudate Cluster 1 | 0.11 | 0.03 | < 0.001*** |
| Caudate Cluster 2 | 0.02 | 0.03 | 0.54 |
| Putamen Cluster 1 | 0.13 | 0.03 | < 0.001*** |
| Putamen Cluster 2 | 0.13 | 0.03 | < 0.001*** |
| Drinking phase (After) | 0.16 | 0.03 | < 0.001*** |
| Drinking phase (Never) | 0.04 | 0.05 | 0.41 |
| Nacc Cluster 2*Drinking phase (After) | -0.17 | 0.03 | < 0.001*** |
| Caudate Cluster 1*Drinking phase (After) | -0.19 | 0.03 | < 0.001*** |
| Caudate Cluster 2*Drinking phase (After) | -0.02 | 0.03 | 0.58 |
| Putamen Cluster 1*Drinking phase (After) | -0.22 | 0.03 | < 0.001*** |
| Putamen Cluster 2*Drinking phase (After) | -0.22 | 0.03 | < 0.001*** |
| Nacc Cluster 2*Drinking phase (Never) | 0.02 | 0.05 | 0.72 |
| Caudate Cluster 1*Drinking phase (Never) | 0.03 | 0.05 | 0.58 |
| Caudate Cluster 2*Drinking phase (Never) | -0.04 | 0.05 | 0.41 |
| Putamen Cluster 1*Drinking phase (Never) | 0.03 | 0.05 | 0.55 |
| Putamen Cluster 2*Drinking phase (Never) | 0.03 | 0.05 | 0.55 |
| **Approx. significance of smooth terms** |  |  |  |
|  | **edf** | ***F*** | ***p*** |
| Age | 3.45 | 12.61 | < 0.001*** |
| **Random effects** |  |  |  |
|  | ***sd*** | **95% CI** | |
| Participant (Intercept) | 0.30 | [0.27, 0.33] | |
| Age (Slope) | 0.012 | [0.010, 0.014] | |
| Residuals | 0.67 | [0.66, 0.68] | |

*Note.* Edf = effective degrees of freedom. * = *p* < 0.05, ** = *p* < 0.01, *** = *p* < 0.001.

**Table S4**. Post-hoc pairwise comparisons results for alcohol frequency.

| **NAcc Cluster 1** | | | | |
| --- | --- | --- | --- | --- |
| **Contrast** | **Estimate** | **S.E.** | ***t*** | ***p*** |
| Before – After | -0.16 | 0.03 | -5.91 | < 0.001* |
| Before – Never | -0.04 | 0.05 | -0.83 | 0.41 |
| After – Never | 0.12 | 0.05 | 2.56 | 0.01 |
| **NAcc Cluster 2** | | | | |
| **Contrast** | **Estimate** | **S.E.** | ***t*** | ***p*** |
| Before – After | 0.01 | 0.03 | 0.48 | 0.63 |
| Before – Never | -0.06 | 0.05 | -1.18 | 0.24 |
| After – Never | -0.07 | 0.05 | -1.50 | 0.13 |
| **Caudate Cluster 1** | | | | |
| **Contrast** | **Estimate** | **S.E.** | ***t*** | ***p*** |
| Before – After | 0.03 | 0.03 | 1.06 | 0.29 |
| Before – Never | -0.07 | 0.05 | -1.37 | 0.17 |
| After – Never | -0.09 | 0.05 | -2.03 | 0.04 |
| **Caudate Cluster 2** | | | | |
| **Contrast** | **Estimate** | **S.E.** | ***t*** | ***p*** |
| Before – After | -0.14 | 0.03 | -5.20 | < 0.001* |
| Before – Never | -0.001 | 0.05 | -0.02 | 0.98 |
| After – Never | 0.14 | 0.05 | 2.99 | 0.05 |
| **Putamen Cluster 1** | | | | |
| **Contrast** | **Estimate** | **S.E.** | ***t*** | ***p*** |
| Before – After | 0.06 | 0.03 | 2.07 | 0.04 |
| Before – Never | -0.07 | 0.05 | -1.41 | 0.16 |
| After – Never | -0.12 | 0.05 | -2.66 | 0.01 |
| **Putamen Cluster 2** | | | | |
| **Contrast** | **Estimate** | **S.E.** | ***t*** | ***p*** |
| Before – After | 0.06 | 0.03 | 2.35 | 0.02 |
| Before – Never | -0.07 | 0.05 | -1.42 | 0.16 |
| After – Never | -0.13 | 0.05 | -2.84 | 0.004 |

*Note.* Edf = effective degrees of freedom.

* *p* is significant at Bonferroni-adjusted α of 0.002.

**Table S5**. Exploratory developmental trajectories of striatal seed-based functional connectivity based on significant Clusters from the alcohol initiation analysis.

| **Seed-Cluster** | | | | | | |
| --- | --- | --- | --- | --- | --- | --- |
|  | **Parametric coefficients** | | | | | |
| **NAcc Cluster 1** |  | **Estimate** | **S.E.** | | ***p*** | |
|  | Intercept | -0.11 | 0.03 | | < 0.001* | |
|  | Sex (Male) | 0.20 | 0.03 | | < 0.001* | |
|  | Drinking Group (Initiated Drinking) | 0.01 | 0.03 | | 0.65 | |
|  | **Approx. significance of smooth terms** | | | | | |
|  |  | **edf** | ***F*** | | ***p*** | |
|  | Age | 1 | 4.03 | | 0.04 | |
|  | Age*Drinking Group (Initiated Drinking) | 3.24 | 10.78 | | < 0.001*** | |
|  | **Random effects** | | | | | |
|  |  | ***sd*** | **95% CI** | | | |
|  | Participant (Intercept) | 0.29 | [0.22, 0.38] | | | |
|  | Age (Slope) | 0.007 | [0.003, 0.02] | | | |
|  | Residuals | 0.589 | [0.57, 0.60] | | | |
|  | **Parametric coefficients** | | | | | |
| **Caudate Cluster 2** |  | **Estimate** | **S.E.** | ***p*** | | |
|  | Intercept | -0.07 | 0.03 | 0.02* | | |
|  | Sex (Male) | 0.13 | 0.03 | < 0.001* | | |
|  | Drinking Group (Initiated Drinking) | 0.02 | 0.03 | 0.62 | | |
|  | **Approx. significance of smooth terms** | | | | | |
|  |  | **edf** | ***F*** | ***p*** | | |
|  | Age | 1 | 7.15 | 0.01* | | |
|  | Age*Drinking Group (Initiated Drinking) | 1 | 0.03 | 0.86 | | |
|  | **Random effects** |  |  | | | |
|  |  | ***sd*** | **95% CI** | | | |
|  | Participant (Intercept) | 0.35 | [0.32, 0.38] | | | |
|  | Age (Slope) | 0.00003 | n.s. | | | |
|  | Residuals | 0.62 | [0.61, 0.64] | | | |
| **Putamen Cluster 1** | **Parametric coefficients** | | | | | |
|  |  | **Estimate** | **S.E.** | | | ***p*** |
|  | Intercept | -0.07 | 0.03 | | | 0.02 |
|  | Sex (Male) | 0.20 | 0.04 | | | < 0.001* |
|  | Drinking Group (Initiated Drinking) | -0.04 | 0.04 | | | 0.24 |
|  | **Approx. significance of smooth terms** | | | | | |
|  |  | **edf** | ***F*** | | | ***p*** |
|  | Age | 1 | 13.30 | | | < 0.001* |
|  | Age*Drinking Group (Initiated Drinking) | 1 | 1.21 | | | 0.27 |
|  | **Random effects** | | | | | |
|  |  | ***sd*** | **95% CI** | | | |
|  | Participant (Intercept) | 0.36 | [0.28, 0.46] | | | |
|  | Age (Slope) | 0.01 | [0.006, 0.02] | | | |
|  | Residuals | 0.62 | [0.61, 0.64] | | | |

*Note.* Edf = effective degrees of freedom. * *p* is significant at Bonferroni-adjusted α of 0.016.

**Table S6.** Association between seed-based functional connectivity and frequency of alcohol use.

| **Seed-Cluster** | **Parametric coefficients** | | | | |
| --- | --- | --- | --- | --- | --- |
| **NAcc Cluster 1** |  | Estimate | S.E. | *t* | *p* |
|  | Intercept | 19.16 | 1.26 | 15.23 | < 0.001* |
|  | Sex (Male) | 3.13 | 1.76 | 1.78 | 0.08 |
|  | **Approx. significance of smooth terms** | | | | |
|  |  | edf | *F* | *p* |  |
|  | Age | 3.92 | 422.39 | < 0.001* |  |
|  | Age*Nacc Cluster 1 | 2 | 3.14 | 0.044 |  |
| **NAcc Cluster 2** | **Parametric coefficients** | | | | |
|  |  | Estimate | S.E. | *t* | *P* |
|  | Intercept | 19.19 | 1.26 | 15.29 | < 0.001* |
|  | Sex (Male) | 2.91 | 1.76 | 1.65 | 0.10 |
|  | **Approx. significance of smooth terms** | | | | |
|  |  | edf | *F* | *P* |  |
|  | Age | 3.91 | 421.03 | < 0.001* |  |
|  | Age*Nacc Cluster 2 | 2.35 | 0.38 | 0.82 |  |
| **Caudate Cluster 1** | **Parametric coefficients** | | | | |
|  |  | Estimate | S.E. | *t* | *P* |
|  | Intercept | 19.21 | 1.26 | 15.28 | < 0.001* |
|  | Sex (Male) | 3.05 | 1.76 | 1.73 | 0.08 |
|  | **Approx. significance of smooth terms** | | | | |
|  |  | edf | *F* | *P* |  |
|  | Age | 3.92 | 420.35 | < 0.001* |  |
|  | Age*Caudate Cluster 1 | 2 | 1.38 | 0.25 |  |
| **Caudate Cluster 2** | **Parametric coefficients** | | | | |
|  |  | Estimate | S.E. | *t* | *P* |
|  | Intercept | 19.13 | 1.26 | 15.22 | < 0.001* |
|  | Sex (Male) | 2.88 | 1.76 | 1.64 | 0.10 |
|  | **Approx. significance of smooth terms** | | | | |
|  |  | edf | *F* | *P* |  |
|  | Age | 3.91 | 417.85 | < 0.001* |  |
|  | Age*Caudate Cluster 2 | 2 | 1.70 | 0.18 |  |
| **Putamen Cluster 1** | **Parametric coefficients** | | | | |
|  |  | Estimate | S.E. | *t* | *P* |
|  | Intercept | 19.39 | 1.26 | 15.48 | < 0.001* |
|  | Sex (Male) | 2.92 | 1.76 | 1.66 | 0.10 |
|  | **Approx. significance of smooth terms** | | | | |
|  |  | edf | *F* | *P* |  |
|  | Age | 3.91 | 427.40 | < 0.001* |  |
|  | Age*Putamen Cluster 1 | 3.37 | 4.42 | 0.005* |  |
| **Putamen Cluster 2** | **Parametric coefficients** | | | | |
|  |  | Estimate | S.E. | *t* | *p* |
|  | Intercept | 19.34 | 1.26 | 15.41 | < 0.001* |
|  | Sex (Male) | 2.97 | 1.76 | 1.69 | 0.09 |
|  | **Approx. significance of smooth terms** | | | | |
|  |  | edf | *F* | *p* |  |
|  | Age | 3.91 | 426.81 | < 0.001* |  |
|  | Age*Putamen Cluster 2 | 2.83 | 4.29 | 0.01 |  |

*Note.* Edf = effective degrees of freedom. * *p* is significant at Bonferroni-adjusted α of 0.008.

**Supplemental Figures**

**K-means clustering results**

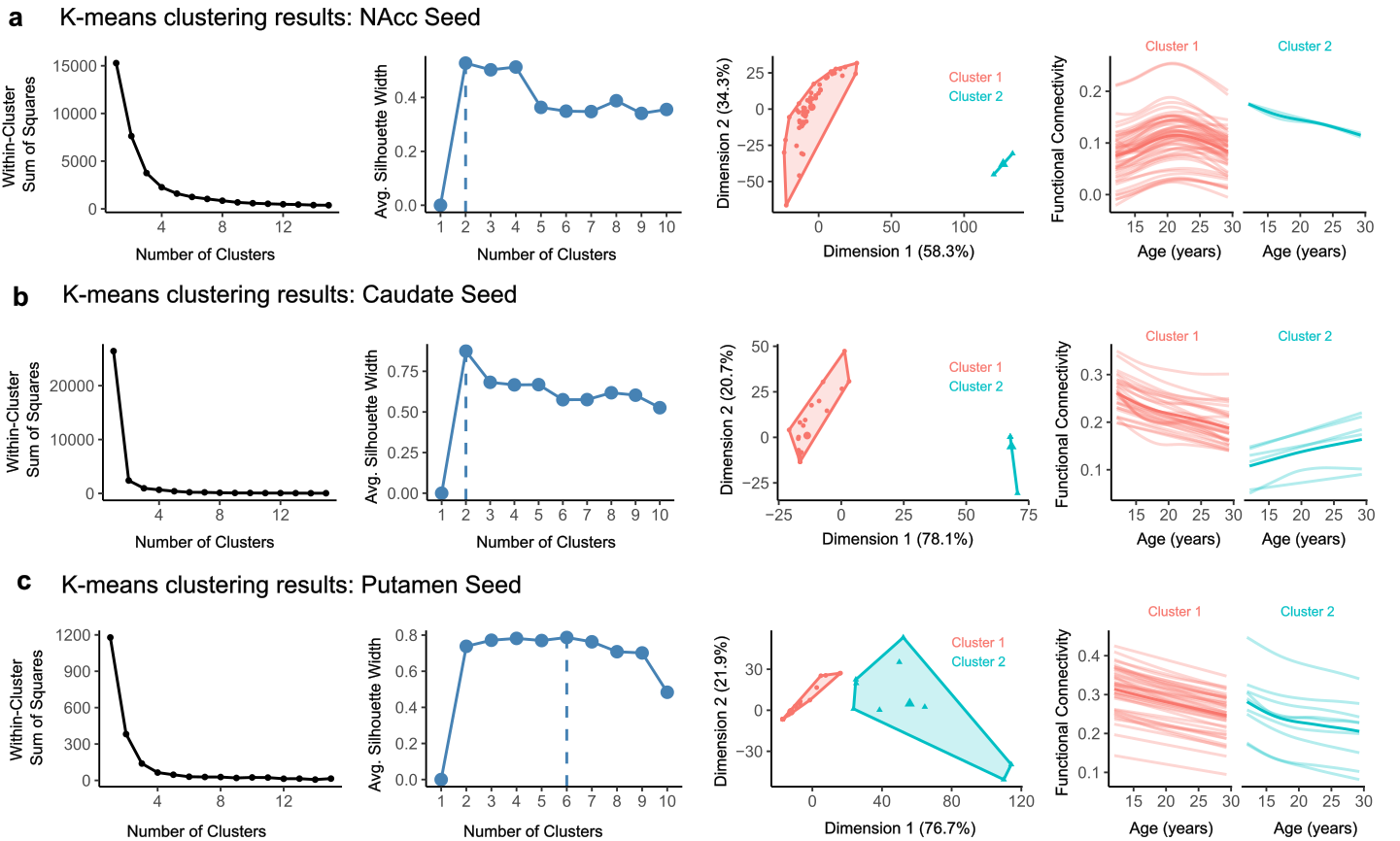

**Figure S1.** K‑means clustering results for three seed regions **(a–c)**. Each row shows clustering diagnostics and resulting cluster structure for: **(a)** NAcc seed, **(b)** Caudate seed, and **(c)** Putamen seed. Within each row, the first panel displays the elbow plot illustrating the within‑cluster sum of squares across values of k. The second panel shows the average silhouette width for each k, with the selected solution indicated by the dashed vertical line. The third panel presents a two‑dimensional projection of the final clustering solution, with points colored by cluster and polygons outlining cluster boundaries. The fourth panel shows age‑related functional connectivity trajectories for each cluster, with individual trajectories plotted alongside their corresponding cluster centroid.

**Supplemental Alcohol use initation results**

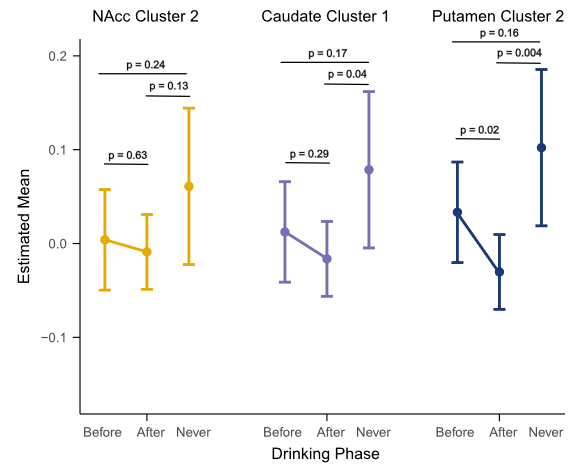

**Figure S2**. Estimated mean functional connectivity by alcohol‑initiation phase across clusters derived from NAcc, Caudate, and Putamen seed‑based analyses. Each panel displays the estimated marginal means (± 95% CI) for participants classified as initiating alcohol use before, after, or never during the study period. Separate panels correspond to Cluster 1 and Cluster 2 for each seed region (NAcc, Caudate, Putamen). Horizontal brackets indicate pairwise comparisons among drinking‑phase groups, with associated p‑values shown above each comparison. Patterns illustrate how functional connectivity differs across drinking‑initiation phases within each cluster.

**Supplemental Drinking phase difference smooths results**

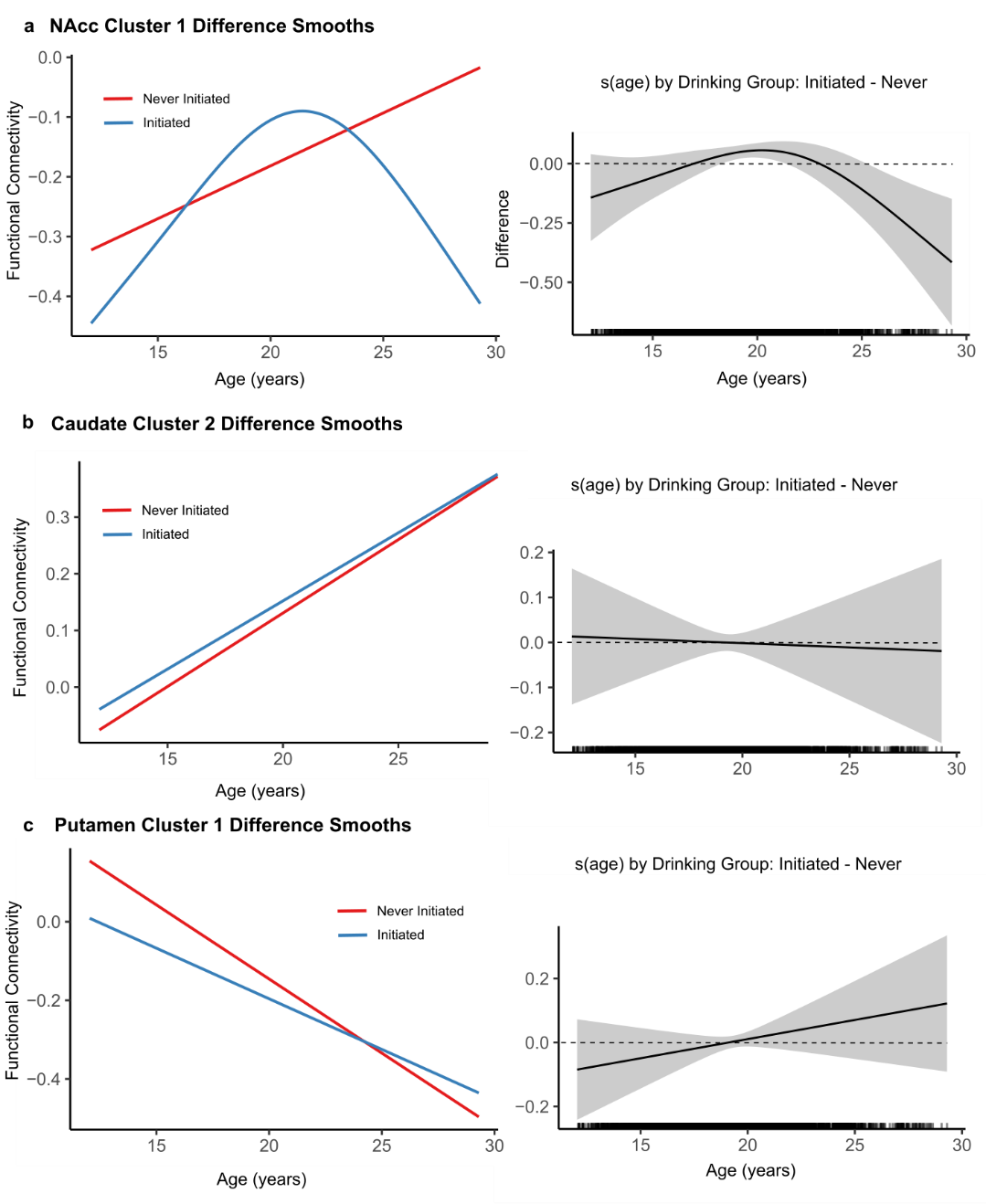

**Figure S3.** Functional connectivity trajectories by drinking‑initiation status for clusters derived from NAcc **(a)**, Caudate **(b)**, and Putamen **(c)** seed‑based analyses. In each row, the left panel shows smooth age‑related functional connectivity trajectories for participants who never initiated alcohol use (red line) and those who initiated (blue line). The right panel depicts the corresponding difference smooth (Initiated – Never), with 95% confidence intervals shown in gray. The horizontal dashed line at zero indicates no group difference; confidence intervals that exclude zero identify age periods where the trajectories differ significantly between drinking‑initiation groups.

**Supplemental time-varying effects models examining the effects of FC on days of alcohol use in the past year**

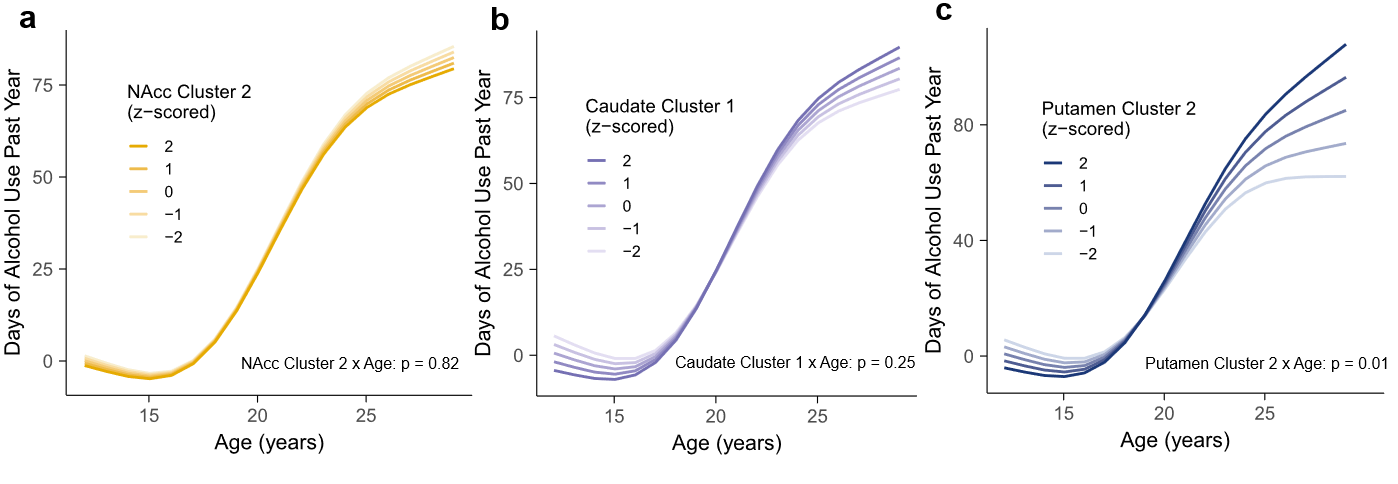

**Figure S4.** Time‑varying effect model (TVEM) results depicting the association between age and past‑year days of alcohol use across functional connectivity clusters derived from NAcc Cluster 2 **(a)**, Caudate Cluster 1 **(b)**, and Putamen Cluster 2 **(c)** seeds. Within each panel, curves represent predicted days of alcohol use across adolescence and emerging adulthood for individuals at different standardized values (z‑scores) of functional connectivity within each cluster. Together, the panels illustrate how the strength and developmental timing of the association between functional connectivity and alcohol use varies across clusters and seed regions. A Bonferroni correction was applied for six models, resulting in an adjusted significance threshold of p = 0.008.

**Supplemental analyses examining the effects of FC on days on binge drinking alcohol use in the past year**

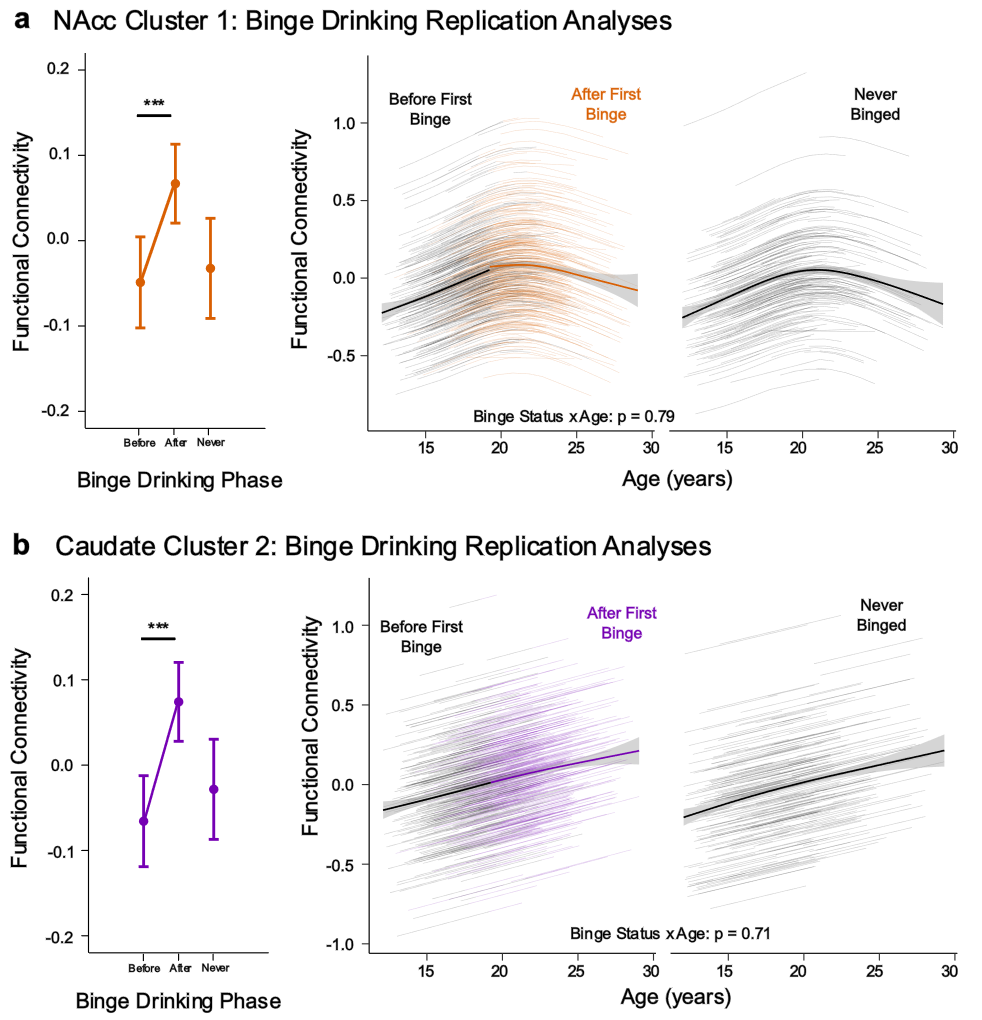

**Figure S5**. Relationship between corticostriatal FC and the initiation of binge-drinking behavior. *** = p < 0.001. **(a left)** Estimated marginal means (± 95% CI) for NAcc Cluster 1 FC in participants classified as initiating binge drinking before, after, or never during the study period. Horizontal brackets indicate pairwise comparisons among binge drinking phase groups. **(a right)** NAcc Cluster 1 FC trajectories based on whether an individual initiated binge drinking or if they never initiated binge drinking. **(b)** Caudate Cluster 2 results. Panels parallel those in (a).

**Supplemental analyses examining the effects of FC on days of nicotine use in the past month**

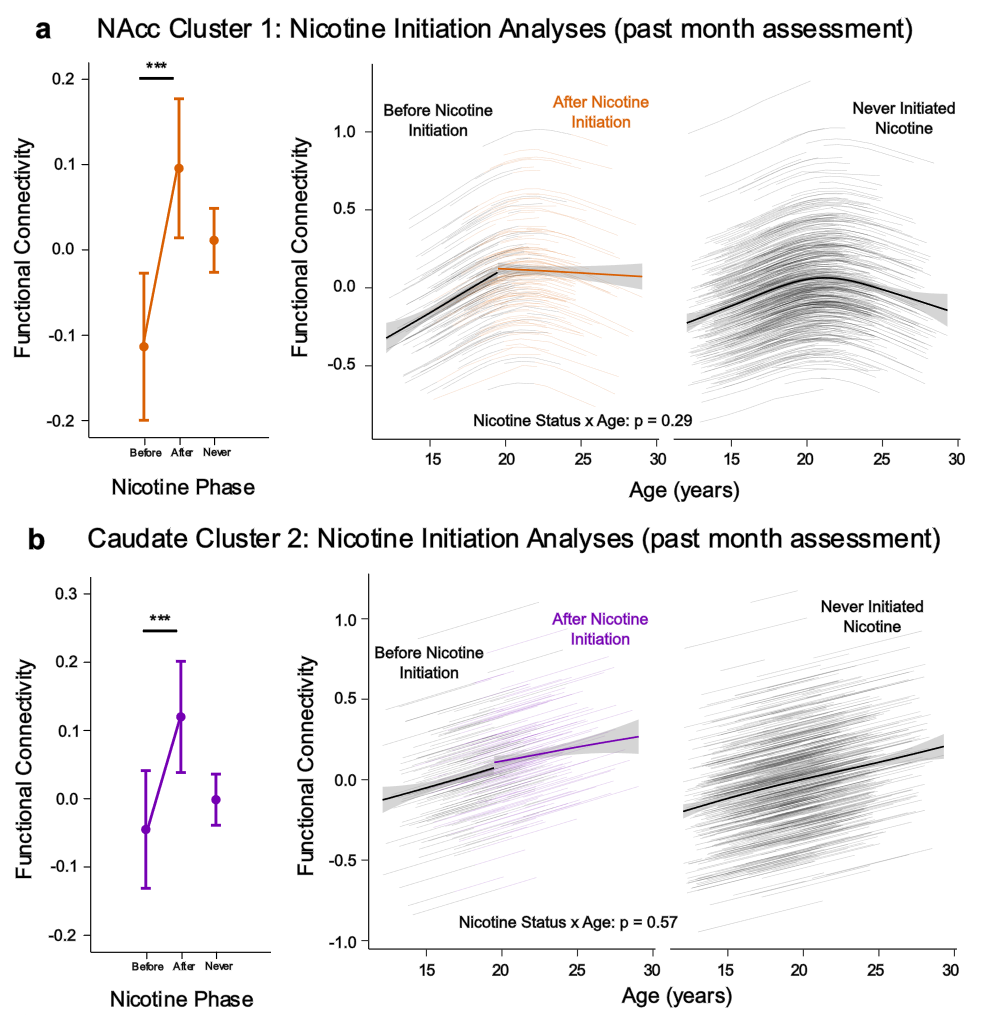

**Figure S6**. Relationship between corticostriatal functional connectivity and initiation of nicotine, based on retrospective self‑report of past‑month use at each visit. *** = p < 0.001. **(a left)** Estimated marginal means (± 95% CI) for NAcc Cluster 1 FC in participants classified as initiating nicotine use before, after, or never during the study period. Horizontal brackets indicate pairwise comparisons among nicotine phase groups. **(a right)** NAcc Cluster 1 FC trajectories based on whether an individual initiated nicotine use or if they never initiated nicotine use. **(b)** Caudate Cluster 2 results. Panels parallel those in (a).

**Supplemental analyses examining the effects of FC on days of cannabis use in the past month**

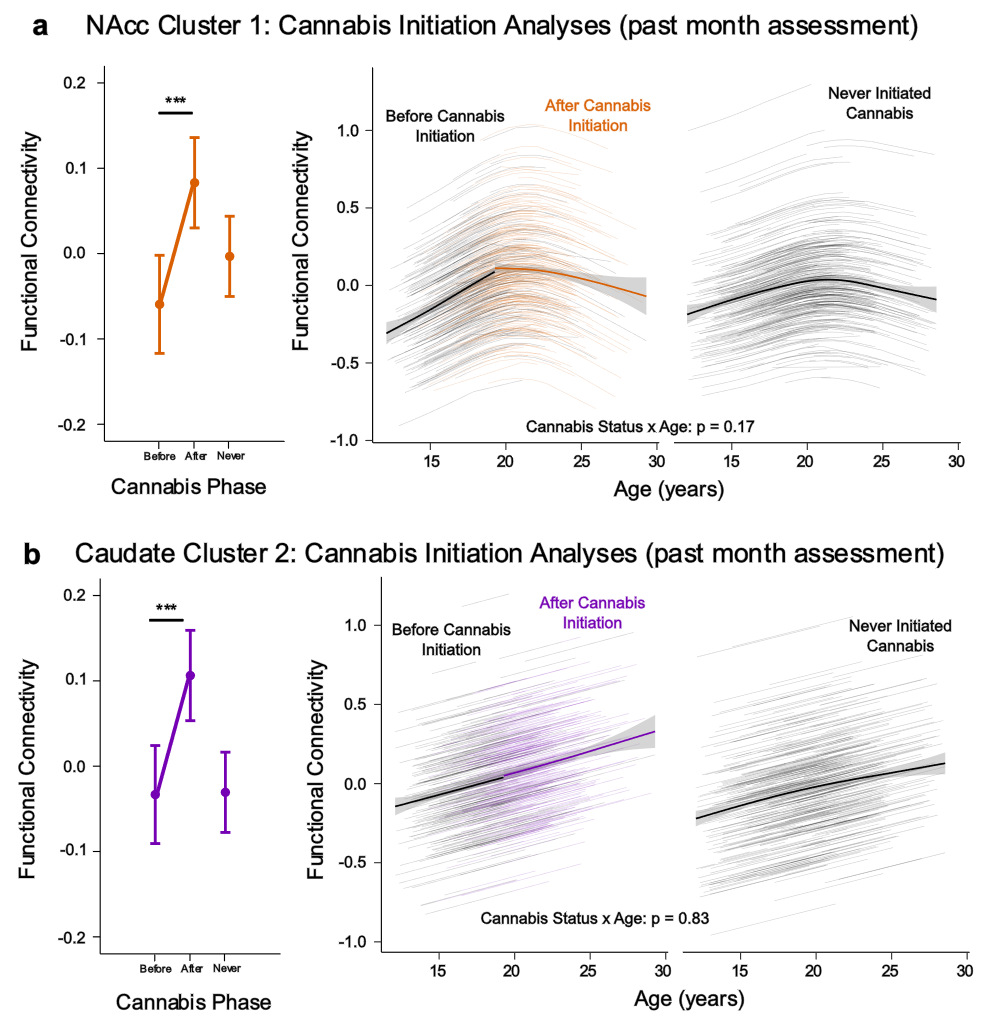

**Figure S7**. Relationship between corticostriatal functional connectivity and initiation of cannabis use, based on retrospective self‑report of past‑month use at each visit. *** = p < 0.001. **(a left)** Estimated marginal means (± 95% CI) for NAcc Cluster 1 FC in participants classified as initiating cannabis use before, after, or never during the study period. Horizontal brackets indicate pairwise comparisons among cannabis use phase groups. **(a right)** NAcc Cluster 1 FC trajectories based on whether an individual initiated cannabis or if they never initiated cannabis use. **(b)** Caudate Cluster 2 results. Panels parallel those in (a).

**Supplemental analyses examining the effects of FC on days of cannabis use in the past year**

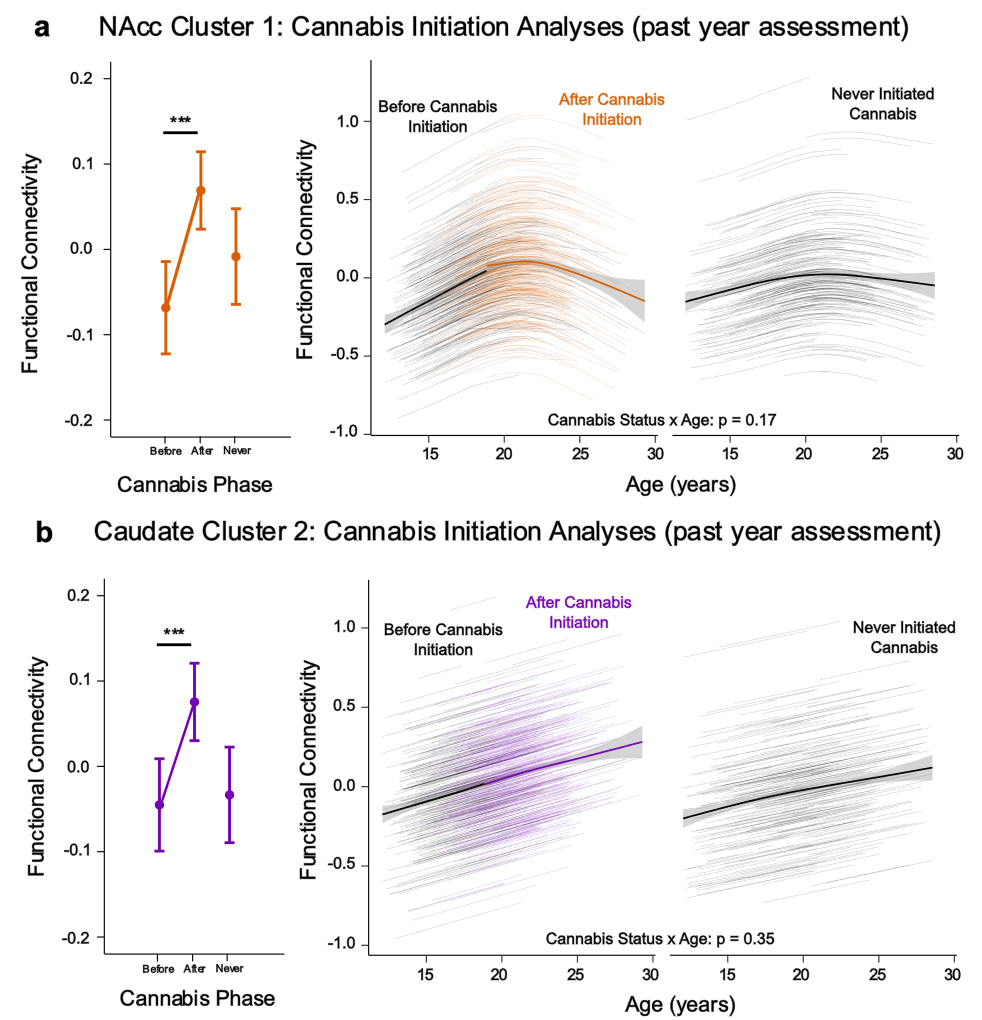

**Figure S8**. Relationship between corticostriatal functional connectivity and initiation of cannabis use, based on retrospective self‑report of past‑year use at each visit. *** = p < 0.001. **(a left)** Estimated marginal means (± 95% CI) for NAcc Cluster 1 FC in participants classified as initiating cannabis use before, after, or never during the study period. Horizontal brackets indicate pairwise comparisons among cannabis use phase groups. **(a right)** NAcc Cluster 1 FC trajectories based on whether an individual initiated cannabis or if they never initiated cannabis use. **(b)** Caudate Cluster 2 results. Panels parallel those in (a).
